# The mutational landscape of *EGFR-*, *MYC-*, and *Kras-* driven genetically-engineered mouse models of lung adenocarcinoma

**DOI:** 10.1101/048058

**Authors:** David G. McFadden, Katerina Politi, Arjun Bhutkar, Frances K. Chen, Xiaoling Song, Mono Pirun, Philip M. Santiago, Caroline Kim, James T. Platt, Emily Lee, Emily Hodges, Adam P. Rosebrock, Roderick Bronson, Nicholas D. Socci, Gregory Hannon, Tyler Jacks, Harold Varmus

## Abstract

Genetically-engineered mouse models (GEMMs) of cancer are increasingly being utilized to assess putative driver mutations identified by large scale sequencing of human cancer genomes. In order to accurately interpret experiments that introduce additional mutations, an understanding of the somatic genetic profile and evolution of GEMM tumors is necessary. Here, we performed whole exome sequencing of tumors from three GEMMs of lung adenocarcinoma driven by mutant EGFR, mutant Kras or by overexpression of MYC. Tumors from EGFR- and Kras- driven models exhibited respectively 0.02 and 0.07 non-synonymous mutations/megabase, a dramatically lower average mutational frequency than observed in human lung adenocarcinomas. Tumors from models driven by strong cancer drivers (mutant EGFR and Kras) harbored few mutations in known cancer genes, whereas tumors driven by *MYC*, a weaker initiating oncogene in the murine lung, acquired recurrent clonal oncogenic *Kras* mutations. In addition, although EGFR- and Kras- driven models both exhibited recurrent whole chromosome DNA copy number alterations, the specific chromosomes altered by gain or loss were different in each model. These data demonstrate that GEMM tumors exhibit relatively simple somatic genotypes compared to human cancers of a similar type, making these autochthonous model systems useful for additive engineering approaches to assess the potential of novel mutations on tumorigenesis, cancer progression, and drug sensitivity.

## INTRODUCTION

Lung cancer remains the leading cause of cancer death worldwide, estimated to have caused 158,000 deaths in the US in 2015 (seer.cancer.gov). Lung adenocarcinoma is the most common form of lung cancer, in both smokers and nonsmokers. Tobacco mutagens cause a high mutation frequency in the somatic genomes of smoking-associated tumors, complicating identification of the genetic drivers of tumor progression (1, 2). An increasing number of somatic alterations that can be targeted by existing drugs or by drug candidates have been identified in lung adenocarcinoma, and several of these agents have demonstrated efficacy in patients (3).

*KRAS* and *EGFR* are the most frequently mutated oncogenes in human lung adenocarcinoma (1, 2, 4, 5). *KRAS* mutations are associated with a strong history of cigarette smoking, whereas *EGFR* mutations are the most frequent oncogene alterations in lung cancers from never-smokers (6). Our groups and others have generated genetically engineered mouse models of *EGFR-* and *KRAS-* mutant lung adenocarcinoma (7-10). These models recapitulate key features of the human disease, including histologic architecture and response and resistance to conventional and targeted therapies (11, 12). Although individual mice develop multifocal disease, only a subset of the primary Kras tumors progress to metastatic disease. A distinct gene expression signature has been shown to distinguish metastatic from nonmetastatic primary tumors in a Kras^G12D^-driven GEMM, suggesting that acquired genetic or epigenetic alterations underlie metastatic progression (13).

Somatic genome evolution in tumors produced in GEMMs remains incompletely characterized. Several studies have described the spectrum of acquired DNA copy number alterations in murine models (14-19). Although these studies reached varying conclusions, it appears that somatic alterations in DNA copy number, especially changes in copy number of certain whole chromosomes, are a common somatic event during tumor evolution in GEMMs. We also recently studied a GEMM of small cell lung cancer using exome and whole genome sequencing. In this model, which is initiated by mutation of the p53 and retinoblastoma (Rb) tumor suppressors, we identified recurrent alterations in the PTEN/PI3K pathway, in addition to previously identified focal DNA amplifications targeting the *Mycl1* oncogene (20-22). Taken together, these studies suggest that, similar to human cancers, GEMM tumors can undergo extensive genome remodeling during tumor progression, and that a subset of these acquired events contributes to cancer progression.

The mutational landscape of carcinogen-induced murine lung adenocarcinomas was also recently described and compared to that in tumors initiated by expression of an oncogenic Kras allele (23). Not surprisingly, single nucleotide mutations, including *Kras* mutations, were more frequently observed in carcinogen-induced tumors. In contrast, secondary DNA copy number alterations were more prevalent in tumors arising in genetically-engineered mice. This further suggests that the path of somatic alteration and selection during tumor progression depends on the specific events that initiate tumorigenesis. As previously described, the carcinogen-treated tumors acquired clonal oncogenic *Kras* mutations. However, it remains unknown whether murine lung adenocarcinomas initiated by other oncogenic drivers, or those harboring combined loss of the tumor suppressor p53, acquire similar or distinct patterns of somatic alteration during tumor evolution and progression. Here, we describe the somatic evolution of a panel of tumors and tumor-derived cell lines derived from genetically-engineered mouse models of lung adenocarcinoma initiated by *Kras*, *EGFR*, and *MYC* (*10*, *24-26*).

## RESULTS

### Mouse models of lung adenocarcinoma acquire few somatic point mutations

We generated a panel of tumor specimens and tumor cell lines from GEMMs of *EGFR-*, *KRAS-*, and *MYC-* mutant lung adenocarcinoma (Figure 1). We performed whole exome sequencing on tumors and cell lines from these models in order to profile the spectrum of genomic alterations acquired during tumorigenesis and progression (Tables 1, S1, S2).

**Figure 1.**
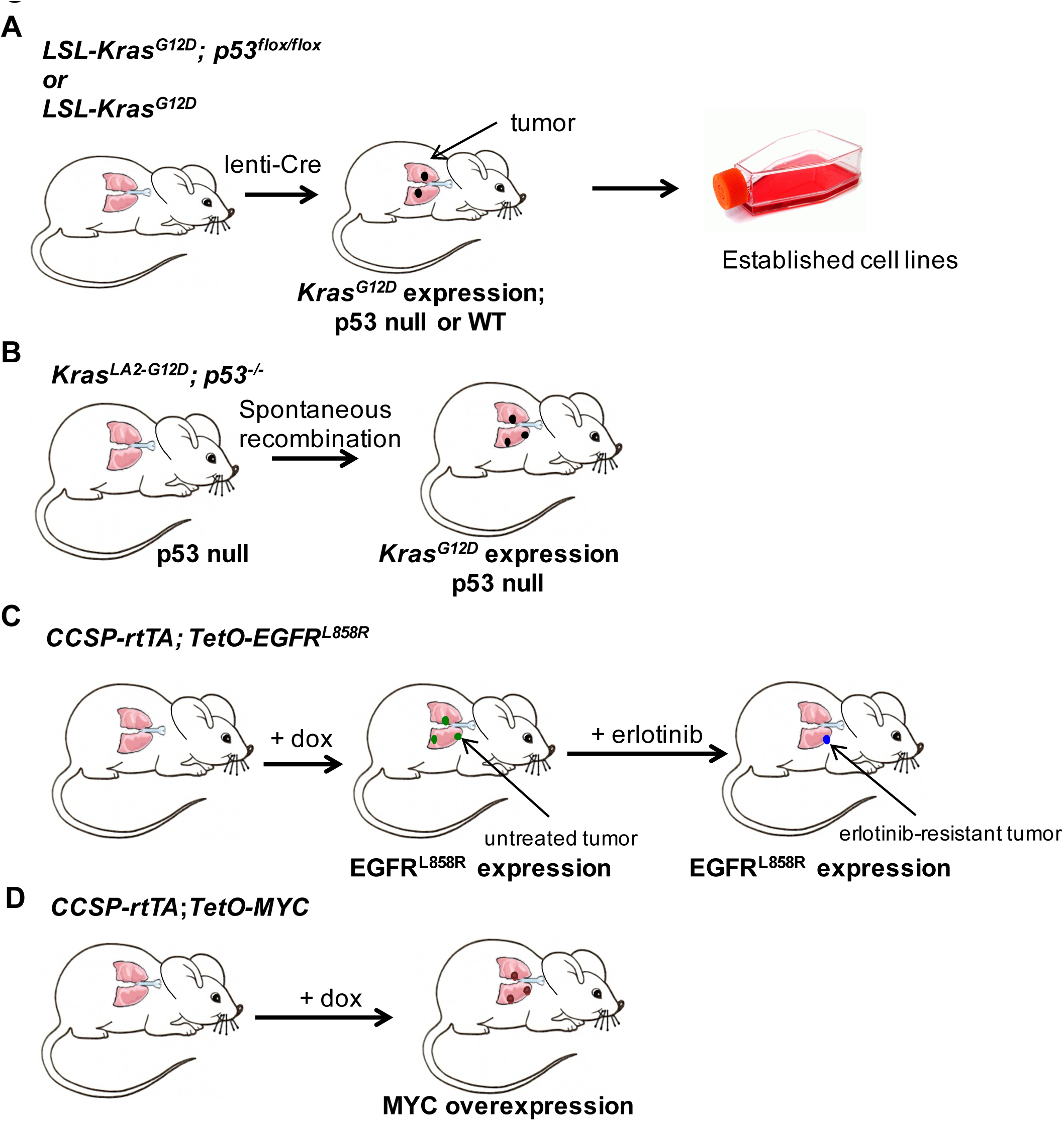
Diagrams illustrating the mouse models of mutant *Kras*, mutant *EGFR* and *MYC-* induced lung adenocarcinoma used in whole exome sequencing. Kras models: A) Mice carrying conditional Kras^LSL-G12D^ and p53^flox/flox^ alleles develop lung adenocarcinomas upon administration of lenti-cre. Cell lines were generated from primary and metastatic lung tumors. Tumors and cell lines were collected for exome sequencing. B) Mice carrying Kras^LA2-G12D^; p53^-/-^ form lung adenocarcinomas spontaneously. Primary tumors were collected for exome sequencing. EGFR model: C) Bi-transgenic *CCSP-rtTA; TetO-EGFR*^*L858R*^ mice were treated with doxycycline at weaning to induce transgene expression (Politi, 2006). Tumors were collected from untreated tumor-bearing mice or mice were treated with erlotinib as described previously until the appearance of resistant tumors (Politi, 2010). Untreated and erlotinib-resistant lung tumors were collected and used for exome sequencing. MYC model: D) Bitransgenic *CCSP-rtTA; TetO-MYC* mice were treated with doxycycline at weaning to induce transgene expression. Overexpression of MYC in type II pneumocyte leads to the development of lung adenocarcinomas that were collected for whole exome sequencing (Tran, PLoS One 2008).

**Table 1:** Lung adenocarcinoma GEMMs utilized for the sequencing study. GEMMs are grouped by initiating driver event, and columns show the tumor type (studied as a cell line or primary tumor tissue), genotype of animals, method of cancer gene induction, and number of samples from each model included in the study. Additional details on the models are found in the Methods and Figure S1.

| Initiating Driver | Cell line or Tumor | Genotype | Tumor Induction | Treatment | Number Studied | Reference |
| --- | --- | --- | --- | --- | --- | --- |
| Kras-G12D | Cell line | Kras-LSL-G12D; Trp53 FL/FL | Lenti-Cre | n/a | 15 | Jackson et al, 2005 |
| Kras-G12D | Tumor | Kras-LSL-G12D; Trp53 FL/FL | Lenti-Cre | n/a | 9 | Jackson et al, 2005 |
| Kras-G12D | Tumor | Kras-LSL-G12D | Lenti-Cre | n/a | 8 | Jackson et al, 2001 |
| Kras-G12D | Tumor | Kras-LA2; Trp53 -/- | spontaneous recombination | n/a | 4 | Johnson et al, 2001 |
| EGFR-L858R | Tumor | CCSP-rtTA; tetO::EGFR-L858R | doxycycline | none | 10 | Politi et al, 2006 |
| EGFR-L858R | Tumor | CCSP-rtTA; tetO::EGFR-L858R | doxycycline | erlotinib-resistant | 6 | Politi et al, 2010 |
| MYC | Tumor | CCSP-rtTA; tetO::MYC | doxycycline | n/a | 5 | Tran et al, 2008 |

We initially focused on somatic point mutations. Several methods of mutation calling have been developed but agreement among the various methods is poor (27); at the beginning of our studies, there was no independent evaluation of which method had optimal sensitivity and specificity. Evidence that aggregating calls from multiple methods improves performance was recently described (23, 28). An added challenge was that the methods were developed to call mutations in human tumors, and many had parameters (e.g. background mutation rate) that were optimized for human samples. Therefore, we created a controlled dataset in order to assess the performance of our variantcalling pipeline in a murine background by simulating mixtures of tumor and normal DNA using exon capture of mixtures of germline DNA from different inbred mouse strains (Figure S1A).

As a first step, we generated exon-capture sequencing libraries with tail DNA from inbred C57BL/6 and 129S1/SvImJ mice. In order to simulate tumor subclonal heterogeneity and contamination with infiltrating non-tumor stromal cells, we serially diluted the 129S1/SvImJ library (mimicking “tumor DNA”) into the C57BL/6 library (mimicking “normal DNA”) (Figure S1A). The starting libraries and mixtures were sequenced to a median average depth of 132x (range from 92×-205×). Somatic mutations were identified using both muTect (v1.1.4) and a somatic mutation caller developed by our group (hereafter referred to as the HaJaVa caller) based on the GATK UnifiedGenotyper with filtering to call somatic events (as described in detail in the Methods sections) (29). Using this approach, individual germline polymorphisms can be traced through the serially diluted libraries, mimicking somatic variant detection in tumors. This dataset was used to estimate the false positive and false negative rates at decreasing allelic fraction.

At the indicated depth of coverage, muTect was highly sensitive, particularly at low allelic fractions that might be found in samples with a low fraction of tumor cells or as a result of subclonal mutational events, but it exhibited a higher false positive rate. In contrast, the HaJaVa caller exhibited a higher true-positive rate, but also a higher false negative rate at low allelic fraction (Supplemental methods, Figure S1B-D, Table S3). We found that the intersection of the two callers exhibited a lower false-positive rate than either caller alone, reducing missed calls by approximately 50%, with a minimal increase in the false negative rate. Therefore, we used the intersection of the HaJaVa and muTect calling algorithms to identify somatic mutations and to compare datasets among the EGFR, MYC and Kras models.

### Kras-driven models of lung adenocarcinoma

We generated DNA from a large panel of tumors and tumor-derived cell lines from *Kras*^*LSL-G12D*^-based mouse models for whole exome sequencing (Table 1). We sequenced DNA from fifteen tumor cell lines derived from tumor-bearing *Kras*^*LSL-G12D*^*; Trp53*^*fl/fl*^ mice, following the lentivirus-based delivery of cre recombinase (Figure 1)(30). In nine cases, we also sequenced DNA from the parental tumor from which the cell lines were derived. In addition, to determine whether expression of cre recombinase generated unexpected mutations, we sequenced DNA from four tumors arising in the *Kras*^*LA2-G12D*^*;Trp5S*^*-/-*^ model, in which spontaneous recombination at the *Kras* locus, rather than cre-induced recombination, initiated tumorigenesis (31). Finally, in order to assess the potential impact of p53 loss on mutation frequency, we sequenced DNA from eight tumors initiated in *Kras*^*LSL-G12D*^*; Trp53*^*WT*^animals.

We observed a median nonsynonymous mutation frequency of 0.07/Mb (including missense and nonsense mutations and mutations affecting splicing signals; Figure 2A, range 0.00-0.46) in Kras-driven tumors (cell lines are excluded from this analysis – see below). Interestingly, we did not observe a statistically different mutation frequencies between *Trp53* null (0.07 mutations/Mb) and wild-type tumors (0.06 mutations/Mb, Figure 2B, P value=0.60). Tumors initiated in the *Kras*^*LSL-G12D*^*; Trp5*^*fl/fl*^ model harbored numbers of mutations (0.09 mutations/Mb) similar to those in the *Kras*^*LA2-G12D*^*;Trp53*^*-/-*^ model (0.03) mutations/Mb, P value=0.1, Figure 2C). With the *Kras*^*LSL-G12D*^*; Trp53*^*fl/fl*^ model, we observed a statistically significant increase in the mutation frequency in tumor-derived cell lines (0.25 mutations/Mb) compared to primary tumor specimens (0.07 mutations/Mb) (Figure 2D, p-value=0.001). This may reflect the presence of subclonal mutations present in the genomes of cell line-founder clones that would be expected to be enriched during the generation of tumor cell lines. We did not observe a predilection for specific base transitions or transversions in the tumors or cell lines (Figure S2).

**Figure 2.**
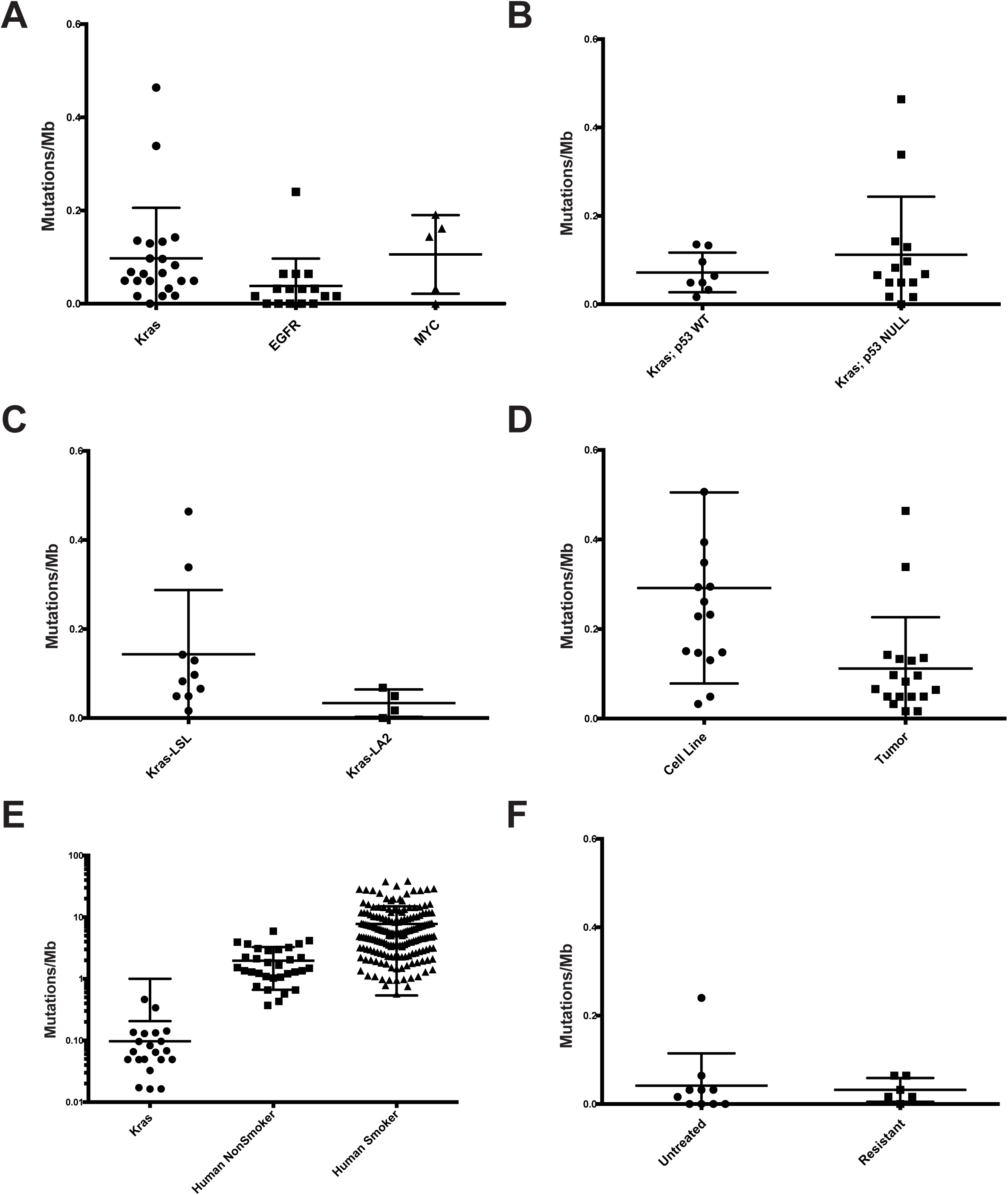
Low mutational burden in GEMM models of lung cancer. Dot plots showing the non-synonymous mutation frequency observed from WES datasets in murine LUADs induced by oncogenic Kras (tumors from either the LA2 or LSL models; tumor-derived cell lines are excluded, see below), EGFR or overexpression of MYC (A), *Trp53* null vs. wild-type tumors (B), *Kras*^*LA2-G12D*^*;p53*^*-/-*^ vs. *Kras*^*LSL-G12D*^*; Trp53*^*mi*^ tumors (C), *Kras*^*LSL-G12D*^*; Trp53*^*mi*^ tumors vs. tumorderived cell lines (D), comparison of Kras^G12D^-induced tumors to human lung adenocarcinomas (Ref. 4) from smoking and nonsmoking patients, shown in log scale (E), untreated vs. drugresistant EGFR^L858R^-induced LUADs (F).

We compared these datasets to available sequencing data from human lung adenocarcinoma (Figure 2E). We observed a significantly fewer nonsynonymous mutations in GEMM models than in either smoker-or nonsmoker-associated human lung adenocarcinomas (Kras GEMM, 0.07 mutations/Mb, nonsmoker-associated, 1.97 mutations/Mb, p-value <0.0001, and smoker associated, 7.76 mutations/Mb).

We identified independent recurrent mutations in a number of genes, including *C5ar1*, *Dnahc5, Nyap2, Pcdh15, Pclo, Rngtt, Stil, Tenm4, and Xirp2*. Seven genes (*Csmd1,Hmcn1, Kmt2c, Pcdh15, Pclo, Ttn, Xirp2*) mutated in the mouse Kras^G12D^-induced cell lines and tumors were also recurrently mutated in >15% of samples in human lung adenocarcinomas (LUAD), as reported by The Cancer Genome Atlas (TCGA). Therefore, *Pcdh15, Pclo, Ttn and Xirp2* were recurrently mutated in both the KP mouse model and human lung adenocarcinoma (4)(Figure 3, denoted with an asterisk).

**Figure 3.**
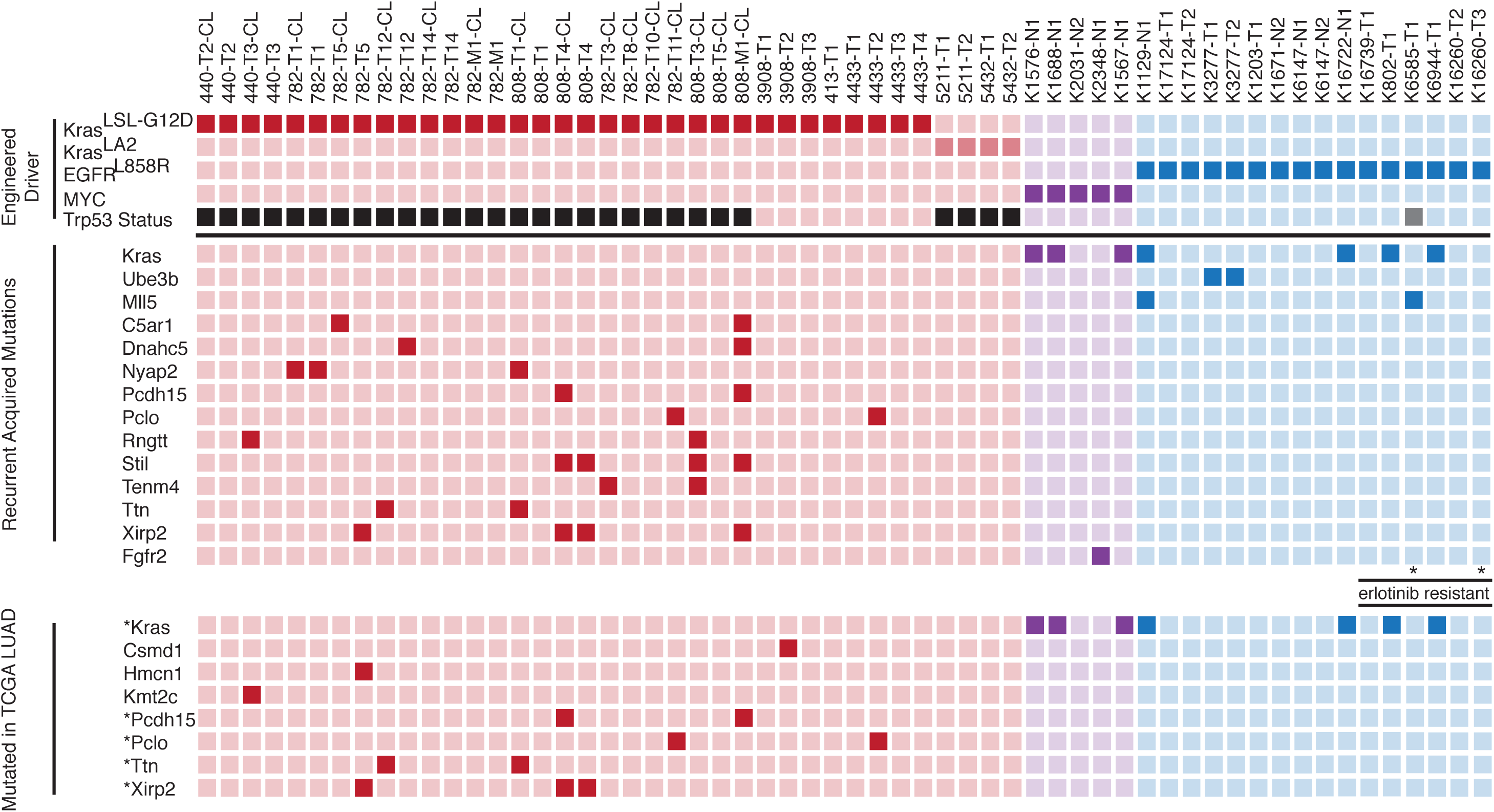
Mutational landscape of oncogene-induced mouse lung adenocarcinomas. Schematic diagram of genes mutated in the mouse lung adenocarcinomas and cell lines. (Top) Kras-, EGFR and MYC-induced tumors are indicated and shaded in red, blue and purple, respectively. The *Trp53* status is indicated (black=null, grey= heterozygous). (Middle) Genes mutated in 2 or more samples are indicated. (Bottom) Genes mutated in the murine tumors that are also mutated in >15% of lung adenocarcinomas analyzed in the TCGA. Note that Csmd1, HmCn1, and Kml2c were not recurrently mutated in murine tumor, whereas asterisk indicated genes recurrently mutated in human and murine tumors. Erlotinib-resistant tumors are indicated. Stars are used to highlight tumors harboring an EGFR T790M mutation (by conventional Sanger sequencing).

As one example, *Xirp2* encodes an actin-binding protein implicated in the maintenance of inner ear hair cell stereocilia and cardiac myocyte remodeling. *Xirp2* was mutated in two independent primary tumors and one pair of primary tumor-metastasis cell lines, the latter of which harbored the same *Xirp2* mutation within a highly conserved region of the Xin actin binding repeats. *Xirp2* has no known role in cancer, yet 21% of human lung adenocarcinomas in the TCGA study harbored mutations in *XIRP2* (4). Review of RNAseq data from the TCGA study revealed very low expression of XIRP2 mRNA in human lung adenocarcinoma, suggesting that mutations in *XIRP2* may be passenger events, despite their high frequency. Alternatively, cells with *XIRP2* mutations might have been selected at an early stage of tumorigenesis but they may not be advantageous during outgrowth of the dominant tumor subclone prior to clinical detection.

We also observed recurrent mutation of *Pclo*, which has been shown to be important for axonal guidance during central nervous system development. *PCLO*, which was mutated in 21% of lung adenocarcinomas in the TCGA study, was also recently identified as a recurrently mutated gene in liver cancers exhibiting a biliary phenotype (32). Knockdown of PCLO RNA in human liver cancer cells led to an increase in cell migration. We identified two mutations in *Pclo*, a nonsense mutation (E577X) and E1850K; the latter resides in a conserved region of the protein with unknown function. Further investigation of the role of Pclo alterations, especially in the context of *Kras* mutations, seems warranted.

Manual review of mutations occurring in a single tumor revealed mutations in several genes encoding regulators of transcription, including several factors involved in chromatin modification and regulation. Among these are mutations in *Brd4* (H965P, within a proline-rich domain of unknown function), *Chd7* (a nonsense mutation, R977X), *Chd8* (H2198R), *Mll3* (T1798S), *Mlxip* (V453G), *Smarcb1(M27R), Smyd4* (H769R), and *Tet1* (C1784X) (Table S4). None of these specific mutations has been identified in human cancers. Therefore, it is difficult to determine if any of these represent driver events. However, mutations in many epigenetic regulators in human tumors are not clustered into “hot spots,” so it is premature to conclude that these are passenger mutations in the mouse model.

### EGFR-driven model of lung adenocarcinoma

Tetracycline-inducible expression of the EGFR^L858R^ mutant in the lung epithelium of transgenic mice leads to the formation of lung adenocarcinomas with bronchioalveolar carcinoma features that are sensitive to treatment with EGFR tyrosine kinase inhibitors like erlotinib (10). Long-term intermittent dosing of the mice with erlotinib leads to the emergence of drug-resistant tumors that harbor some of the molecular features of TKI-resistant human tumors, including a secondary mutation in EGFR, EGFR^T790M^ (12). To determine the mutational load of these tumors, compared to tumors initiated by alternate oncogenes, and to seek genetic differences between untreated and erlotinib-resistant tumors, we performed whole exome sequencing of DNA from ten TKI-naïve and six erlotinib-resistant EGFR^L858R^-induced mouse lung adenocarcinomas. We observed a lower mutational burden in EGFR mutant lung adenocarcinomas compared to Kras-driven tumors (0.02 vs. 0.07 mutations/Mb, P value=0.002)(Figure 2A). Erlotinib-resistant tumors exhibited no difference in mutation frequency compared to untreated EGFR mutant tumors (0.02 vs. 0.02, P value=0.49, Figure 2F). The mutational signature present in the EGFR^L858R^-induced lung adenocarcinomas exhibits a preponderance of C>T transitions, consistent with findings in EGFR mutant human lung adenocarcinomas and all adenocarcinomas from never-smokers (4) (Figure S2).

Recurrent mutations in the tumors were found in *Kras* (n=4, 2 G12V and 2 Q61R), *Mll5* (n=2) and *Ube3b* (n=2). We previously described an oncogenic *Kras* mutation in an erlotinib-resistant murine tumor; however, such mutations have not been described in patients (12). Interestingly, two of the *Kras* mutations observed here (with non reference allele fractions of 0. 38-0.55) were found in the ten tumors not treated with TKI’s, suggesting that these can arise during tumor development independent of treatment. A recent report shows that co-expression of mutant EGFR and KRAS in the same human lung tumor cells can be toxic (33). It is possible that detection of mutations in both oncogenes in some untreated tumors indicates that expression of one of the oncogenes has been down-regulated in at least some tumor cells; in other tumors with both mutations, inhibition of the EGFR kinase activity with a TKI may have been a permissive feature.

*Mll5* (*Kmt2e*), a histone lysine methyltransferase involved in chromatin remodeling, was found to be mutated in two EGFR-induced tumors. Both of the mutations (in an untreated and an erlotinib-resistant tumor) were in disordered domains of the protein and their functional consequences are unknown. *Ube3b*, an E3 Ubiquitin ligase, was mutated in two individual tumors from a single untreated mouse. The same variant was found in both tumors, suggesting that the two tumors are clonally related. According to TCGA reports, *Mll5* and *Ube3b* are altered in 6% and 4% of human LUADs respectively; however, there is no indication at this point of a functional relationship between EGFR mutations and alterations in these genes.

Since a major mechanism of resistance to EGFR inhibitors is a secondary mutation in *EGFR* (*EGFR*^*T790M*^), we examined the whole exome sequencing data to determine whether we could detect reads corresponding to human *EGFR* (since the transgene encodes human *EGFR*). Indeed, we unequivocally detected the *EGFR*^*T790M*^ mutation in one erlotinib-resistant tumor that we had previously shown to harbor this mutation. However, we cannot exclude inadequate depth of sequencing of the human EGFR transgene as an explanation of our failure to identify other cases of secondary T790M mutations of EGFR in these tumors, especially since the exon capture probes target mouse sequences.

### MYC model of lung adenocarcinoma

The low mutation rate observed in the *Kras* and EGFR-induced lung tumors prompted us to hypothesize that tumors induced by strong oncogenic lung drivers might have a lower mutation burden than tumors induced by a less potent lung oncogene. We therefore performed whole exome sequencing of DNA from five lung adenocarcinomas that arose in a mouse model initiated by overexpression of wild type human *MYC*, which has been shown to be a less potent oncogene than mutant *KRAS* or *EGFR* in the murine lung (25). MYC-induced tumors also exhibited a low mutation frequency, comparable to that observed in Kras-induced mouse lung adenocarcinomas (0.14 vs 0.07 mutations/Mb, P value=0.57 Figure 2A). The mutations found in the MYC-induced tumors included oncogenic *Kras* mutations in three of five tumors and an oncogenic mutation in *Fgfr2* (Fgfr2 K659M) in one tumor. Mutations at this residue in *FGFR2* have been shown to activate the intrinsic protein-tyrosine kinase and to cooperate with *MYC* in tumorigenesis (34, 35). The identification of known cancer driver mutations in 4 of 5 MYC-driven tumors is consistent with the suggestion that MYC acts as a less potent tumor initiator that mutant Kras or EGFR in the murine lung.

### Acquired whole chromosome copy number changes are common in mouse lung adenocarcinomas

Given the low point mutation rates observed in the mouse tumors in our GEMM models, we asked whether the tumors harbored alterations in chromosomal or subchromosomal copy numbers. In order to examine somatic changes in DNA copy number in the GEMM tumors, we analyzed the datasets from whole exome sequencing using validated computational methods (36, 37). We first examined the *Trp53* locus, which was anticipated to be deleted in the Kras/p53^fl/fl^ tumors, when Cre recombinase was expressed to initiate tumorigenesis (24). We detected deletion of exons 2-10 in all tumor cell lines derived from those animals, suggesting that the method based on sequence data accurately identified small regions of deletion (Figure S3).

When these methods were applied to the complete set of exome sequencing data, we primarily detected putative whole chromosome gains and losses in the murine lung adenocarcinomas (Figure 4), consistent with prior studies of cancers arising in GEMM’s (16, 17). Manual review of putative focal amplifications and deletions revealed that many were unlikely to be true events. In particular, one signature observed was a set of breakpoints with the same start and stop positions and in tumors that were matched to the same normal sample. These are likely to be artifacts of variable coverage from sequencing of the normal (tail) DNA. Another finding included regions with nearly balanced amplifications and deletions with overlapping annotated duplication regions in the UCSC browser. These are likely to be copy number polymorphisms (Table S5).

**Figure 4:**
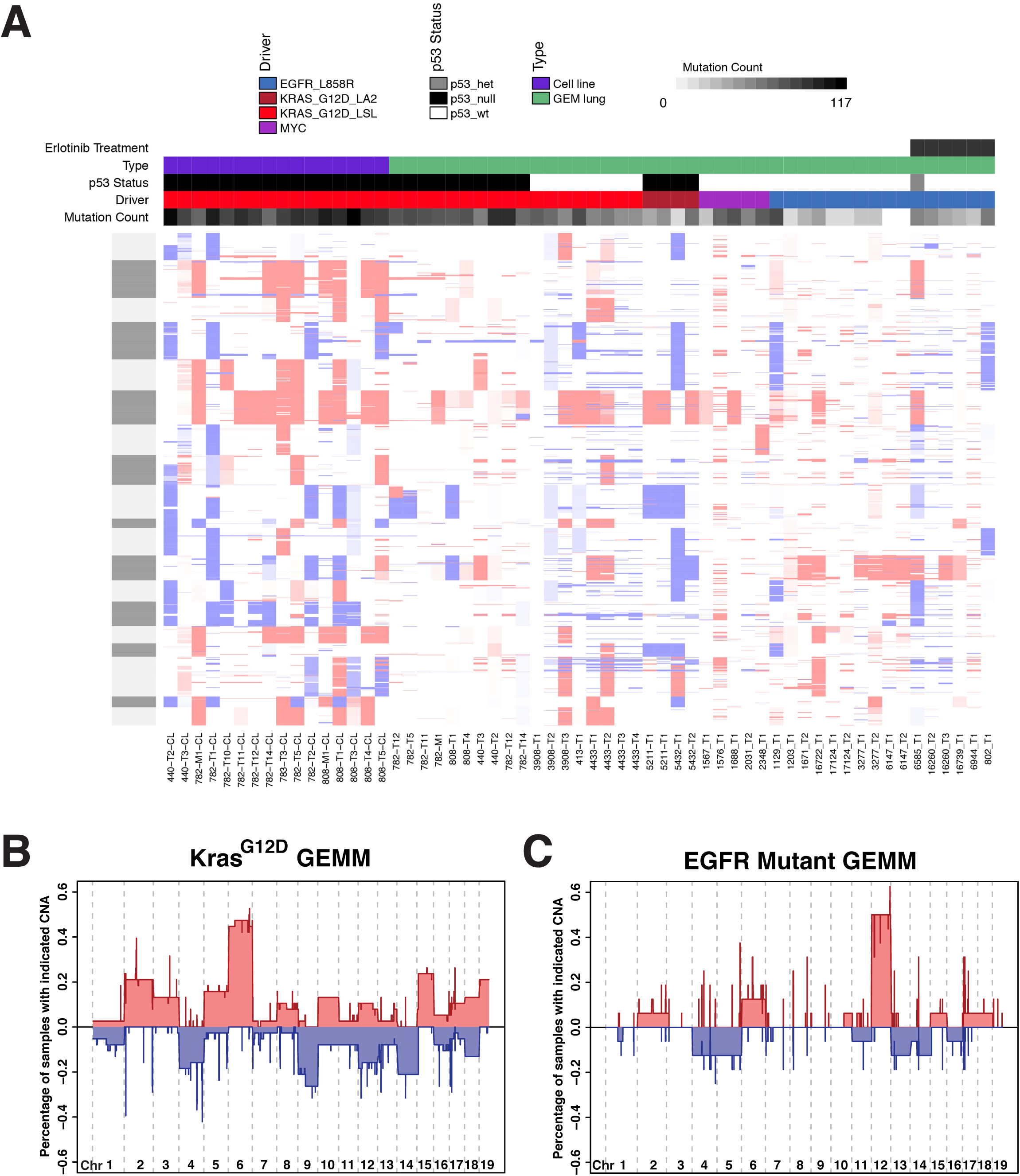
Distinct patterns of DNA copy number alterations in *Kras-* and *EGFR-* driven GEMMs. A) Heat map of DNA copy number alterations across all samples. Red = DNA copy number amplification, blue = DNA copy number loss. Models are grouped by initiating driver allele. Also shown is p53 status, and whether the sample was a tumor or tumor-derived cell line. Point mutation frequency is shown in grey-black box with darker shade representing a higher mutation frequency. B) Recurrent whole chromosome DNA copy number gains in *Kras*^G12D^-driven GEMM tumors and cell lines. C) Recurrent whole chromosome DNA copy number gains in EGFR-driven GEMM tumors.

Although the models we have studied all produce one histological tumor type, lung adenocarcinoma, the tumors that develop in each GEMM show a distinct pattern of recurrent DNA copy number gain or loss (Figure 4A-C, Table S5). Kras-driven tumors and cell lines harbored recurrent gain of Chr6, consistent with prior studies of Kras-induced lung tumors (16, 17, 23, 38). In addition to extra copies of Chr6, we observed recurrent whole chromosome amplification of Chr2, Chr15, and Chr19 (in ≥ 20% of samples), and whole chromosome loss of Chr9 and Chr14 (Table S5). Since Chr6 carries the *Kras* locus, we determined the allelic fraction with the G12D mutation; this analysis suggested that the chromosome with the engineered G12D mutant, not the chromosome with the wild type allele, was responsible for the gain in chromosome number (Figure S4). Similarly, the proto-oncogene *Myc* is on Chr 15, and gain of copies of Chr 15 is the second most frequent whole chromosome alteration observed in this model and consistent with previous work suggesting that Myc function may be necessary for tumor maintenance in Kras-driven GEMMs (39).

Recurrent whole chromosome DNA copy number changes appeared to be less frequent in the EGFR-driven GEMM (Figure 4, Table S6) and a different pattern of changes was observed. For unexplained reasons, an increased number of copies of Chr12 was the most recurrent alteration. It is possible that the unmapped *TetO-EGFR* transgene is integrated in Chr12; gain in copy number might then increase signaling from the *EGFR* oncogene.

We did not observe an anti-correlation between the frequency of point mutations and the fraction of the genome affected by DNA copy number alterations, as previously described (23). This might in part be due to the overall low mutational frequency observed in these tumors, which is approximately an order of magnitude lower than observed in carcinogen-induced models (mean of 185 mutations in urethane-induced and 728 in MNU-induced tumors) (23). Therefore, it is possible that when higher mutation loads are present there is less selection for large-scale DNA copy number alterations across the genome in these models.

## DISCUSSION

Recent improvements in DNA sequencing technologies have spurred the genomic characterization of many types of human cancers, including lung adenocarcinomas. These datasets have revealed much information about the mutational profiles of this cancer, and identified several novel putative oncogenes and tumor suppressors. However, unraveling the complexity of these datasets and distinguishing driver and passenger mutations remain significant challenges, particularly in highly mutated genomes such as smoking-associated lung adenocarcinoma. In contrast, in this report, we have observed a very low mutation frequency in EGFR-, Kras- and MYC-driven GEMM tumors, regardless of tumor genotype. The low mutation frequency in the murine tumors is consistent with prior studies of other GEMMs and suggests that the number of mutations necessary for the development and progression of invasive lung adenocarcinomas in mice is small (19, 40).

We detected recurrent whole chromosome gains and losses in the EGFR and Kras-driven models. These observations raise the question of whether copy number alterations are contributing to tumorigenesis in these models. Considering that we observed recurrent Chr6 amplification, which encodes the engineered *Kras*^*LSL-G12D*^ allele as well as several components of the MAPK signaling pathway, in Kras-driven models, we speculate that cooperating oncogenes and or tumor suppressors located in the regions of whole chromosome gain or loss indeed contribute to tumor progression. However, the identity of the driver events in the amplified and deleted regions remains to be determined. These observations suggest that amplification of the signal from the initiating oncogene may be the most important somatic event in these models to drive tumor progression (38).

Previous work also described changes in gene expression during tumor progression, suggesting that epigenetic alterations contribute to progression in these models (13). The detection of several mutations in transcriptional regulators and chromatin-remodeling factors in our models (see Figure 3) is consistent with these findings, although we have not determined directly whether any of the observed mutations accelerate tumor progression or lead to specific changes in chromatin in these models.

We found evidence for clonal selection during the emergence of drug resistance and during the generation of tumor-derived cell lines. Tumors harvested from mice with *EGFR-* mutant lung cancer harbored oncogenic lesions known to confer primary or acquired resistance to TKIs (for example, *Kras* mutations and the *EGFR* T790M mutation, respectively). In addition, drug-resistant tumors exhibited a higher overall mutation burden compared to untreated samples reflecting increased complexity of these resistant tumors. Similarly, in the Kras-driven model, tumor cell lines harbored a higher mutation frequency compared to the parental tumors, suggesting clonal selection during outgrowth of cell lines.

Despite the low frequency of observed somatic events in the GEMM tumors, each model exhibited distinct features. In contrast to the *Kras* and *EGFR* models, which harbored few mutations known to act as cancer drivers, four out of five *MYC*-induced tumors harbored oncogenic mutations in *Kras* or in *Fgfr2*. The acquisition of potent driver mutations in these tumors suggests *MYC* over-expression sensitizes cells to transformation by cooperating with spontaneous *Kras* or *Fgfr* somatic mutations. In contrast, even in the absence of *Trp53*, Kras-driven tumors did not acquire mutations in known tumor suppressors or oncogenes. This highlights the potency of this oncogene and strongly suggests that the initiating genetically-engineered allele is a critical determinant of acquired events in these models.

The overall non-synonymous mutation burden in human lung adenocarcinomas is 6.86 mutations/Mb (lung TCGA). This is approximately 50-fold higher than the median mutation burden observed in any of the mouse lung adenocarcinomas studied here. In part, this is likely to reflect the lack of carcinogen exposure. However, the mutation rate in never smokers (1.97 mutations/Mb) remains over 10-fold higher than that observed in our lung cancer models (4). The rapid development of tumors in mice may also contribute to the reduced complexity of the cancer genome in these models compared to human lung adenocarcinoma. As previously described, the copy number profiles of mouse tumors were generally characterized by largescale whole chromosome gains or losses (14, 15, 17, 18). In contrast, human tumors exhibited both large-scale and focal amplifications and deletions, perhaps also reflecting differences in carcinogen exposures in tumors in the two species (41).

Our findings have important implications for the optimal use and further development of GEMMs, particularly considering the ease with which these modifications can be generated using new genome editing methods (42-44). Although we have not sequenced to the extreme depth necessary to identify mutations in very small subpopulations of tumor cells, the genomic profile of these models appears to be much less complex than most human cancers. The genomic complexity of lung cancer is at the heart of drug resistance and appears to be an important determinant of the response to immune checkpoint inhibitors (45).

We previously reported that tumors in the *Kras*^*LSL-G12D*^*; Trp53*^*ml*^ model exhibit very modest immune cell infiltrates (46). This is consistent with very few neoantigens generated as a result of the very low number of somatic mutations that arise during tumor development as we describe here. However, expression of a strong T-cell antigen in the model induces a potent T-cell response, which is subsequently suppressed at later stages of tumor progression (46). Therefore, it is important to consider the low mutation frequency exhibited in tumors in these GEMMs when designing therapeutic studies or studies of drug resistance studies. At the same time, the uniformity of the programmed somatic mutations and low acquired mutation frequency observed in tumors in these models are important experimental strengths, making the models well suited to reproducible mechanistic studies and genetic screening. Efforts to model genomic complexity in GEMMs, using mutagens, transposons, or engineered loss of DNA repair pathways are approaches to further optimize GEMMs for studies of sensitivity and resistance to therapies and could identify new drivers of progression and metastasis that cooperate with the initiating engineered mutations.

## METHODS

### Mouse models

All animals studies were performed under approved IACUC protocols at MIT and MSKCC. Tumor induction was performed in *Kras*^*LSL-G12D*^*; p53*^*fl/fl*^ and *Kras*^*LSL-G12D*^ mice as previously described with 2.5 X 10^4^ lentivirus particles per animal (30). Tumors were isolated and tumor-derived cell lines were generated as previously described, for tumor-derived cell lines (13). Histological analysis was performed on a piece of each tumor in order to assess tumor purity and histological subtype of lung cancer. Tumors were induced in the *TetO-EGFR*^*L858R*^ and *TetO-MYC* models by feeding the mice doxycycline impregnated food as previously described (10).

### Exome Sequencing

DNA was purified from tumor tissue and tumor cell lines using standard methods. Sonication of 2ug genomic DNA was performed using a Diagenode Bioruptor, and size selection was performed using dual selection using AMPure beads as previously described (47). Exon capture was performed using Roche SeqCap EZ all-exon mouse kits. Post-capture libraries were sequenced on an Illumina HiSeq instrument.

### Variant Calling

The pipeline to call somatic mutations was predominately composed of standard programs that have been used in human tumor analysis with the addition of a custom caller that was optimized to reduce false positives at the expense of some sensitivity to low allele frequency events. The final mutation list was the intersection of these two calling algorithms in the hope that artifacts given rise to false positives in one would be filtered out by the other.

Raw sequence files were first pre-processed to remove the sequencing adapters. Then clipped reads were mapped to the standard mm9 genome for the Kras model or to mm9+transGene (hEGFR or hMYC) hybrid genomes for the EGFR and MYC models. BWA ALN (version 0.5) was used to make maps. The reads were marked with read groups and sorted, then duplicates were removed using the PICARD toolkit. These initial bam files were post-processed using the standard GATK packages; indel realignment was followed by base quality recalibration. The post-processed bam files were then called using two separate mutation callers: MuTect (v1.1.4) in high confidence mode (HC) and a custom caller built around the GATK Unified Genotyper, with a set of filters to improve specificity. An intersection of these calls were post-filtered for artifacts by removing any events that were in a database of likely germline events (see supplemental for details). The calls were annotated with Annovar and a list of “functional” mutations was created that contained missense, non-sense and splice site mutations.

### DNA Copy Number

Copy number was determined by first computing normalized log ratios between tumors and matched normals. This was done taking by taking the bam files from the mutation-calling pipeline and computing the coverage for each exon target region using bedtools. The raw coverage numbers for each tumor normal pair were normalized with a robust regression method that normalized not only for total depth but also for the local GC content around each target region using the loess function from R. The log (base 2) ratio of T to N was computed from the normalized coverage and this was then segmented using the Circular Binary Segmentation method (CBS) of ref 27. A post segmentation normalization was then used to center the diploid peak at logR = 0. To find segments that were either amplified or deleted we used the RAE algorithm (28), which computes a sample-dependent soft threshold (sigmoid function) for each Tumor/Normal pair based on the noise of that pair. This gives a value of 0-1 for both amplifications and deletions, which roughly indicates fractional amount of each alteration. These values were then average over all samples to give the fraction of each region that was amplified or deleted in a given set of samples.

## ACKNOWLEDGEMENTS

Funding for this work was provided by a grant from the Starr Cancer Consortium to T.J., G.H. and H.V. Additional support was provided by the Howard Hughes Medical Institute (to T.J. and G.H.), by the NIH (NCI K08 160658 to D.G.M., R01CA120247 to K.P. and H.V. R01 CA121210 to K.P., Cancer Center Support Grants P30 CA016359 and P30 CA008748, SPORE grant P50 CA140146-05, and UL1TR000457) and by Uniting Against Lung Cancer (to X.S.). The authors would like to acknowledge Richard Cook and Alla Leshinsky (Biopolymers core facility, KI Swanson Biotechnology Center) and the Hope Babette Chang Histology Facility in the Swanson Biotechnology Center.

